# Bacterioplankton taxa compete for iron along the early spring-summer transition in the Arctic Ocean

**DOI:** 10.1101/2022.02.07.479392

**Authors:** Fernando Puente-Sánchez, Luis Macías, Karley L. Campbell, Marta Royo-Llonch, Vanessa Balagué, Pablo Sánchez, Javier Tamames, C.J. Mundy, Carlos Pedrós-Alió

## Abstract

Microbial assemblages under sea ice of Dease Strait, Canadian Arctic, were sequenced for metagenomes and metatranscriptomes of the small size fraction (0.2 to 3 µm). The community from early March was typical for this season, with *Alpha*- and *Gammaproteobacteria* as the dominant taxa, followed by *Thaumarchaeota* and *Bacteroidetes*. Towards summer, *Bacteroidetes* and particularly the genus *Polaribacter*, became increasingly dominant, followed by the *Gammaproteobacteria*. Analysis of genes responsible for microbial acquisition of iron showed an abundance of ABC transporters for divalent cations and ferrous iron. The most abundant transporters, however, were the outer membrane TonB dependent transporters of iron-siderophore complexes. The abundance of iron acquisition genes suggested this element was essential for the microbial assemblage. Interestingly, *Gammaproteobacteria* were responsible for most of the siderophore synthesis genes. On the contrary, *Bacteroidetes* did not synthesize siderophores but accounted for most of the transporters, suggesting a role as cheaters in the competition for siderophores as public goods. Likely, this cheating ability of the *Bacteroidetes* contributed to their dominance in summer.

## Introduction

Climate change over the recent decades is evident across the Arctic Ocean through the decline in seasonal sea ice coverage [(1), (2), (3)]. This reduction of sea ice impacts the microorganisms living in the ice and underlying water. For example, a shift towards smaller algae has been observed in phytoplankton [(4)]. The distinctive conditions for life in polar environments involve prolonged periods of ice cover and darkness in winter, generally low water temperatures, and extended irradiance in summer [(5)] The transition from winter to spring and summer involves, thus, dramatic changes in the environment and these are reflected in the composition of the microbial assemblages [(6), (7)]. Prokaryotic taxa abundant in winter such as *Thaumarchaeota* or *Deltaproteobacteria* disappear during the spring, while *Alpha* and *Gammaproteobacteria* and in particular *Bacteroidetes* increase in abundance towards the summer [(8)]. However, the traits that these different microbial taxa use to become dominant are unknown. This is partially due to the logistic difficulties of sampling during the early part of the year. Thus, we carried out a time series through the ice melt transition from early March to late July in Dease Strait of the Canadian Arctic.

Iron is an essential nutrient that is scarce in surface waters due to the insolubility of the most common form Fe(III) and to the fact that sources are of terrestrial origin. As a result, between 15 and 40% of the world’s oceans are limited by iron [(9), (10)]. Even in areas where growth is limited by other nutrients (such as the Arctic Ocean) microorganisms need to develop strategies to obtain iron. A common method to get iron in aquatic media is to use siderophores [(11)]. These are small molecular weight compounds that are soluble in water and have a very high affinity for Fe(III). They are synthesized by some microorganisms and usually released in the environment. Then, with the appropriate receptors, the iron-siderophore complexes can be taken up.

Siderophores, however, require a complex set of enzymes to be synthesized and excreted in the medium. And, therefore, they require considerable amount of energy [(12)]. Some bacteria do not synthesize siderophores but have the receptors for them. These bacteria act as cheaters, exploiting “public goods” that they have not contributed to create. Due to these peculiarities, the interactions between siderophore producers and non-producers have been explored with cultures of marine *Vibrio* to illustrate competition for public goods [(13)] and the evolutionary implications of these interactions have been reviewed [(14)]. Most experimental information, however, comes from studies with pure cultures of *Proteobacteria*, particularly of different *Vibrio* and *Pseudomonas* strains. But the issue has not been analyzed in nature where the large diversity of coexisting microorganisms may result in complex interactions.

In the Arctic, *Bacteroidetes* become the most abundant taxon of the assemblage in summer. On the other hand, *Proteobacteria*, that are the most abundant taxa in winter and early spring, decrease in abundance (8). One remarkable difference between *Proteobacteria* and *Bacteroidetes* is the capability to synthesize siderophores. *Proteobacteria* have been shown to synthesize dozens of different siderophores (15), while almost no instances of siderophore synthesis are known among *Bacteroidetes* (16). However, *Bacteroidetes* are known to have receptors for iron-siderophore complexes (17), (18). Therefore, we examine the time evolution of the bacterial assemblage and the presence and abundance of iron-related genes through the ice melting period with the aim to test whether the iron acquisition strategies may have a role in the outcome of the competition among taxa.

## Materials and methods

### Sampling

Samples were collected in Dease Strait, lower Northwest Passage, Nunavut, Canada (69.03°N, 105.33°W; Figure 1). Water depth was 60 m and samples were collected between 7 March and 24 June 2014, as part of the 2014 Ice Covered Ecosystem- CAMbridge Bay Process Study (ICE-CAMPS). During the ICE-CAMPS campaign, Dease Strait was covered by first-year ice (1.8-2.1 m thick) with drifted snow ranging from 5 to < 25 cm in depth, until melt commenced on 8 June. Snow melt caused the ice surface to flood by 16 June (19), followed by subsequent drainage and the formation of melt ponds that persisted until ice break-up on 19 July. One week after ice break-up, the site was sampled a final time in ice-free waters on 30 July aboard the *R/V Martin Bergmann*.

**Figure 1.**
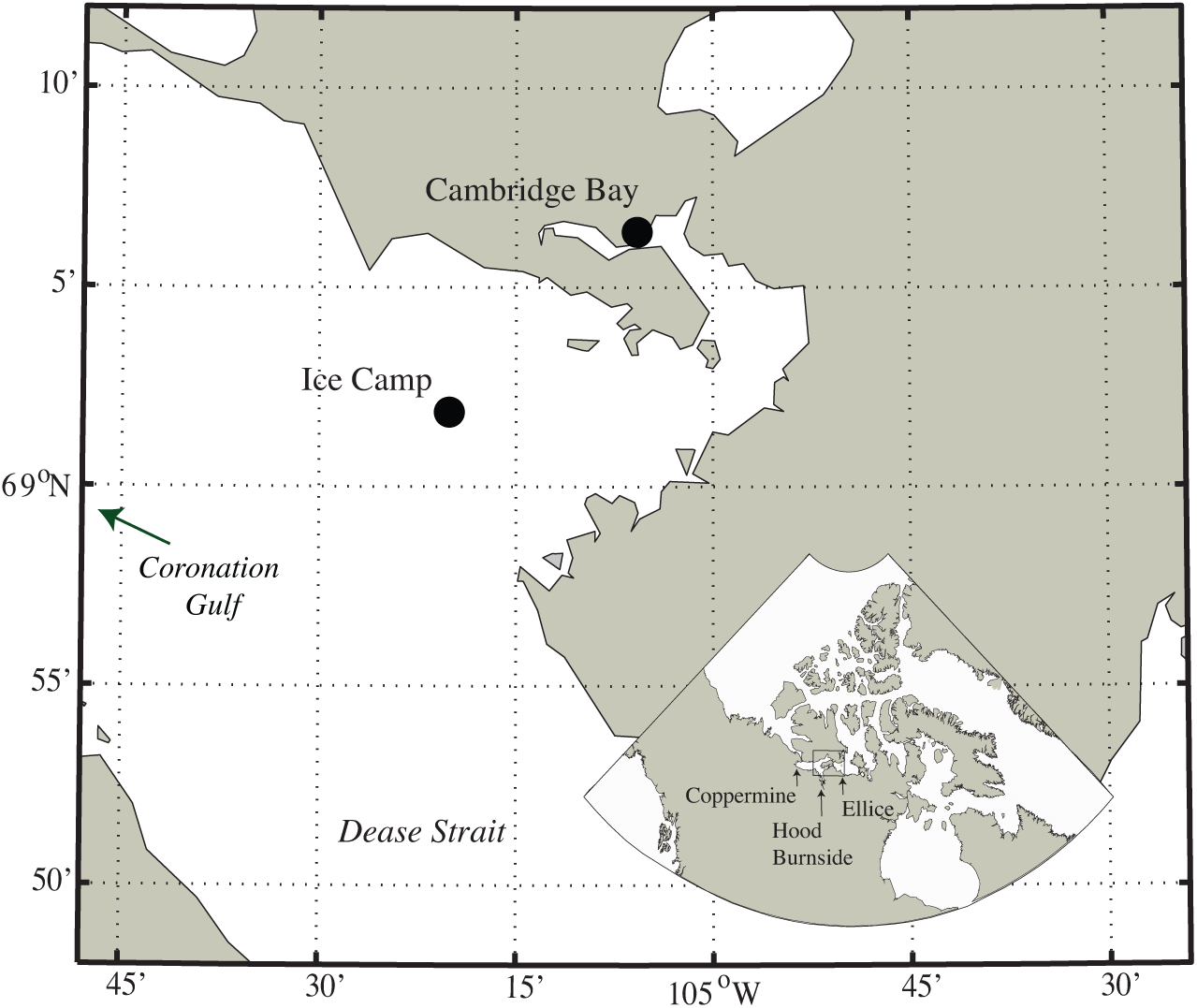
Map of study location near Cambridge Bay, Nunavut. Inset includes approximate locations of the main rivers in the area.

Water samples were collected opportunistically thirteen times, through a 20-cm diameter hole from 7 to 18 March 2014, protected by a portable tent. For the remainder of the ice-covered season (21 April to 24 June 2014), samples were taken from a 2-m^2^ sampling hole maintained within a heated tent. Finally, the 30 July sample was collected, as mentioned, form a ship. A submersible water pump mounted on an under- ice arm was used to collect ice-water interface at 2.5 m. Hydrographic profiles were also taken alongside using a conductivity, temperature and depth probe (RBR XR-620 CTD) equipped with a chlorophyll (chl) *a in vivo* fluorescence sensor. The region is in close proximity to mainland Canada, and receives freshwater inputs from the nearby Coppermine, Hood, Burnside and Ellice rivers (Figure 1). Further details about the study site can be found in Campbell *et al*. (20), (21), (22).

### Physico-chemical measurements

Approximately at 10 m from the location of water sampling, profiles of under-ice photosynthetically active radiation (PAR) were collected between 10:00 am and 12:30 pm local time. This was done using a spherical underwater sensor (LI-193 LI-COR) lowered through an auger hole that took measurements at 1-m depth intervals from the ice-water interface down to 20 m. During measurements the sampling hole was covered to limit contamination by surface radiation. Disturbance of this sites for light measurement was kept to a minimum, where the hole was approached along a single path from the north.

Following transport to laboratory facilities, water samples that were kept in darkness close to, but not below, -2°C, and were immediately processed for chl *a* and nucleic acids. Subsamples for chl *a* were filtered onto GF/F filters (Whatmann) before subsequent pigment extraction in 10 ml of 90% acetone for 24 h in darkness and at 4 °C. Fluorescence was measured before and after acidification with 5% HCl (Turner Designs Trilogy Fluorometer) according to Parsons et al. (1984) and chl *a* concentration was calculated using the equations of Holm-Hansen et al. (23).

In order to collect nucleic acids, approximately 10 l of water was prescreened to remove visibly identifiable plankton and then filtered through a tiered stainless steel large-volume filtration system (Millipore) with a succession of 20, 3 and 0.22 µm filters.

Separate filters were prepared for RNA and DNA. Filters were immediately frozen at -80°C. Here we present data from the 0.22 µm filters only.

### Nucleic acid extraction

DNA was extracted according to the phenol/chloroform protocol detailed by Massana et al. (24). After thawing, frozen filters were cut into small pieces and placed into sterile cryovials. Lysis buffer and lysozyme (final concentration: 1 mg ml^−1^) were added to the cryovials and they were incubated at 37 °C for 45 min. Sodium dodecyl sulfate (10%) and Proteinase K (0.2 mg ml^−1^) were added and the mixture was incubated at 55°C for 1 h. The lysate was mixed twice with an equal volume of phenol/chloroform/isoamyl alcohol (25:24:1, pH 8) and centrifuged at 12 000 × rpm (10 min). A second extraction with chloroform/IAA (24:1) was carried out to remove the residual phenol. The aqueous phase containing the DNA was concentrated and purified by cleaning up with sterile purified water using Amicon Ultra Centrifugal filters (Millipore). The DNA yield and integrity were quantified in a Nanodrop spectrophotometer (NanoDrop 1000 Thermo Fisher) and Qubit fluorimeter (Thermo Fisher). DNA extracts were stored at -80°C.

RNA was extracted after cutting the frozen filters into small pieces and using Qiagen’s RNeasy kit. After a DNase treatment with Ambion’s Turbo DNA-free kit, RNA yield and integrity were quantified with a Nanodrop spectrophotometer (NanoDrop 1000 Thermo Fisher) and a Qubit fluorometer (Thermo Fisher). The RNA extracts were stored at -80°C. Prior to sequencing, ribosomal RNA depletion was performed by Sequentia Biotech (http://www.sequentiabiotech.com/) using Illumina’s TruSeq Stranded Total RNA LT – with RiboZero.

### High-throughput sequencing (HTS)

DNA samples were sequenced in two batches. The first batch included most metagenomic samples and was sequenced at CNAG (https://www.cnag.crg.eu) on the Illumina HiSeq2000 sequencing platform using a TruSeq paired-end cluster kit, v3. The second batch included four metagenomic samples (May 10, June 1, June 15, and July and all metatranscriptomic samples. The numbers of reads per sample are shown in Supplementary Table S1. We used the SqueezeMeta pipeline (v1.3) and the SQMtools R package (25), (26) for all the bioinformatics analyses. Details of the assembly are shown in Supplementary Table S2. Average gene copy numbers per genome were calculated as described in Puente-Sánchez *et al.*, (26) using the median coverage of 10 universal single-copy marker genes (27). These markers are particularly suitable for normalizing metagenomic and metatranscriptomic data to provide estimates of relative per-cell gene copies, because they represent constitutively expressed single-copy housekeeping genes (27). Furthermore, since average copy numbers are normalized against the coverage of well conserved genes, they should be unaffected by the percentage of mapped and/or annotated reads, making them better to compare the abundances of the same function in different samples. Thus, throughout the manuscript we will use average copy numbers per genome when discussing the normalized abundance of individual functions in the microbiome, and percentages of reads assigned to each taxon when discussing taxonomic distributions. Sequences were deposited under NCBI BioProject ID PRJNA803814.

### Taxonomic assignment of ORFs and contigs

For each ORF DIAMOND (v0.9.22) homology searches were performed with its amino acid sequence against the GenBank nr database (downloaded in September, 2019). An e-value cutoff of 1e-03 was set to discard poor hits. The best hit was obtained, and then we selected a range of hits (valid hits) having at least 80% of the bitscore of the best hit, and with an identity to the nr database no smaller than 10% of the identity of the best hit. The Last Common Ancestor (LCA) of all these hits was obtained at diverse taxonomic ranks (from phylum to species). In order to allow for some flexibility, the case of putative transfer events or incorrect annotations in the database, we reported a taxon as the LCA as long as it was supported by 90% of the valid hits.

To ensure trustworthy annotations we also required a minimum identity to the nr database in order to assign ORFs to the different taxonomic ranks. These identity cutoffs were based on Luo *et al.* (28), and were 85, 60, 55, 50, 46, 42 and 40% for the species, genus, family, order, class, phylum and superkingdom ranks, respectively.

Finally, for each contig, we obtained a consensus taxonomy from the annotations of all the ORFs encoded in the contig, such as 50% of all the genes (regardless of whether they could be annotated or not) and 70% of the annotated genes belonging to the same taxon. After mapping of the reads to the contigs using Bowtie2 v2.3.4.1 (29), contig taxonomies were used for calculating global taxon abundances in our samples, while ORF taxonomies were used to calculate the taxonomic distribution of particular functions.

### Functional annotation of open reading frames (ORFs)

The ORFs were functionally annotated into Clusters of Orthologous Groups (COGs) by downloading the eggNOG database (30), selecting the entries annotated as COGs, creating a DIAMOND database and using it for annotation as described in Tamames and Puente-Sánchez (25). COGs were used for functional annotation instead of KEGG orthologs (KOs) or PFAMs, since their finer granularity allowed for a more careful curation of the annotations (see next section).

### Identification of siderophore synthesis genes and pathways

Pathways for siderophore synthesis usually include two types of enzymes. In the first steps, fairly common enzymes are used, such as for example the protein catalyzing the step chorismate to isochorismate. These enzymes are coded by genes that are common, not only to several siderophore synthesis pathways, but also to more general central metabolic pathways. And next, several additional steps that usually require nonribosomal peptide synthetases (NRPs), polyketide synthases, and NRPs-independent siderophore synthetases (NIS) that condense small and large molecules until the complex structures of the siderophores are completed. In many cases, several of these NRPs are common to different pathways and, thus, different gene names are classified in the same COG category. For example, COG3486 has representatives in at least 13 siderophore synthesis pathways (see Supplementary Figure S1). Thus, detecting this COG in metagenomes does not indicate which siderophore pathway is present. On the contrary, some genes are assigned to more than one COG, further complicating the issue. For example, gene *dhbE* in the bacillibactin synthesis pathway is assigned to COGs 0318 (AMP-dependent synthetase and ligase) and COG1021 (2,3- dihydroxybenzoate-AMP ligase). We checked manually all the siderophore synthesis pathways in MetaCyc (31) and added a few more (Supplementary Table S3). We have summarized this complexity in a network (Supplementary Figure S1) that will facilitate identification of both common and unique genes in the synthesis pathways of the different siderophores.

Since identifying the siderophore pathways that are present in metagenomes and expressed in metatranscriptomes is complicated, we adopted the following criteria:

1. In order to state that the pathway for a given siderophore MAY BE present, all the genes in the pathway must be present.
2. In order to state that the pathway IS present, besides presence of all the genes, at least one of the genes must be unique to this pathway, one that is not shared among different pathways.
3. If at least one gene of a pathway is absent, we consider the whole pathway absent.

There was an additional difficulty. A substantial portion of the genes involved in siderophore synthesis belonged to COG0318. This COG includes genes *entE* in salmochelin and enterobactin pathways (three different steps in the pathways), *pvdI* and *pvdL* in pyoverdin (first condensation step), *dhbE* in bacillibactin (two successive steps), *asbC* in petrobactin (a ligase in two steps), and *acsA* in achromobactin (synthase, four steps). It also includes several enzymes unrelated to siderophore synthesis. Thus, we considered the presence of this COG necessary to fulfill criterion 1 above. However, we did not count the reads assigned to this COG in the quantitation of siderophore synthesis genes, because many of them might code for genes in other metabolic routes. We followed a similar procedure for the rest of the COGs, using for quantitation only those which could be unambiguously assigned to siderophore biosynthesis pathways using the high quality – manually curated SwissProt database (32). Some of them, however, should be involved in siderophore synthesis. Therefore, our counts of siderophore synthesis genes are underestimates.

## RESULTS

### Changes in ecosystem parameters

Daylight increased from <12 h during early March to 24 h in 21 May. The consequent increase in surface irradiance was intensified by the progressive melt of surface snow and ice, with PAR transmitted to 2.5 m depth increasing steadily until 12 June, thereafter, increasing nonlinearly (Figure 2a) as melt ponds formed on the surface. Chl *a* increased steadily throughout the study following the increase in PAR, while salinity decreased slightly at the surface due to ice melting (Figure 2b-c).

**Figure 2.**
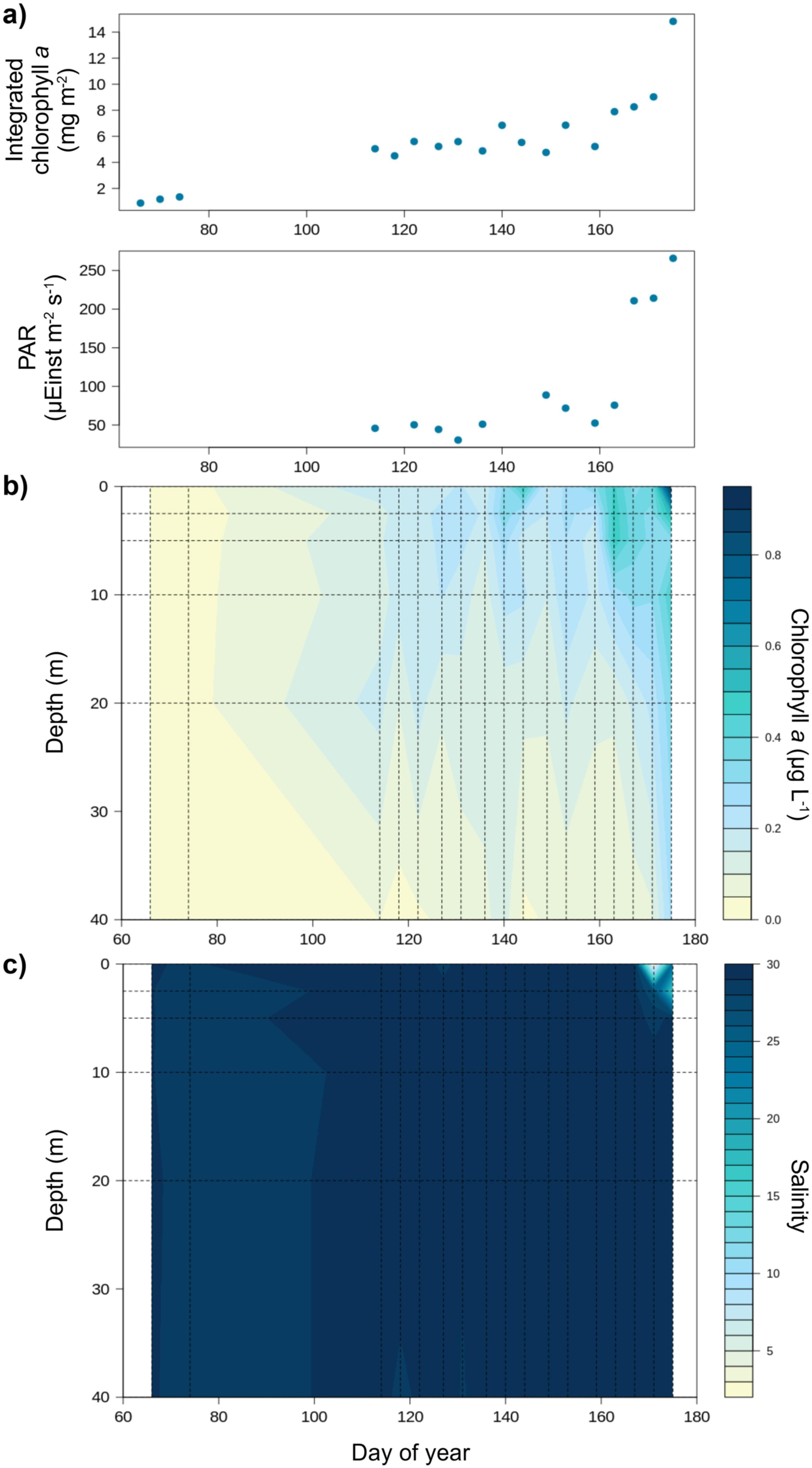
(a) Integrated chlorophyll *a* (mg m^-2^) and PAR (µEinst m^-2^ s^-1^) along day of year. (b) Time-depth diagram of chlorophyll *a* (µg L^-1^). (c) Time-depth diagram of salinity (‰).

### Taxonomic composition of the prokaryotic assemblage

Microbial populations changed along the study (Figure 3a). In early March *Proteobacteria* dominated the assemblage. Specially, *Alphaproteobacteria* accounted for 36% of the reads and *Gammaproteobacteria* for 20%. The two other main groups were *Taumarchaeota* (12%) and *Bacteroidetes* (9%). Other minor components were *Deltaproteobacteria* (5%), *Actinobacteria* (4%), *Betaproteobacteria* (2.5%), *Verrucomicrobia* (2.5%), *Euryarchaeota* (2.5%), *Planctomycetes* (2%), and *Marinimicrobia* (2%). Progressive changes eventually resulted in a different composition in June. For example, on 23 June the main groups were *Bacteroidetes* (55%), *Alphaproteobacteria* (21%), *Gammaproteobacteria* (18%), chloroplasts (12%), and *Actinobacteria* (2%).

**Figure 3.**
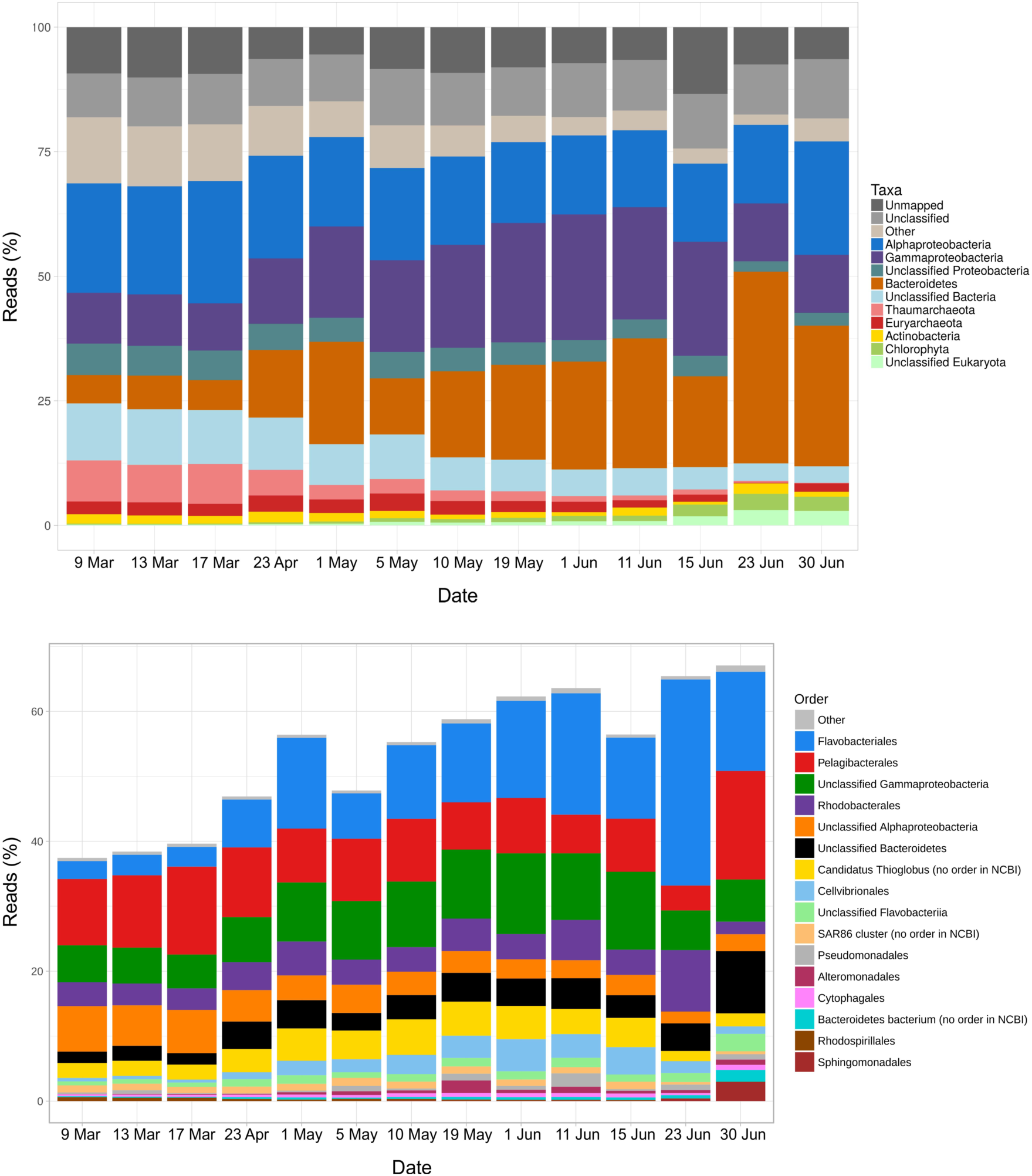
Percent of reads in metagenomes at two different taxonomic levels: phyla and Proteobacteria classes (a) and orders (b).

*Alphaproteobacteria* were, as mentioned, the most abundant group in late winter. The most important orders were *Pelagibacterales* and *Rhodobacterales*, followed by unclassified *Alphaproteobacteria*, and small contributions from *Rhodospirillales* and *Sphingomonadales* (Figure 3b). This is a fairly usual composition in the oceans.

*Gammaproteobacteria* increased up to 1 June, when they reached 37% of the reads and decreased slightly afterwards (Figure 3a). The most abundant group could not be classified beyond the class level. The next taxa in abundance were *Cand*. Thioglobus and *Cellvibrionales* (Figure 3b). Another unusual group was the Order *Pseudomonadales*. In contrast, *Gammaproteobacteria* groups commonly abundant in the oceans such as SAR86 and *Alteromonadales* had only small contributions.

Throughout the study, *Flavobacteriales* were the most abundant *Bacteroidetes* order, followed by unclassified *Bacteroidetes*, unclassified *Flavobacteriia*, and *Cytophagales* (Figure 3b). Most of the increment in *Bacteroidetes* was due to the genus *Polaribacter* (Supplementary Figure S2). In effect, at the beginning of the study, the *Bacteroidetes* reads were fairly well distributed among seven taxa: N55, NS9, *Marinoscillum*, NS4, *Polaribacter*, *Owenweeksia* and *Sufflavibacter* in order of abundance and several other minor components. In late June, on the other hand, 60% of the reads belonged to *Polaribacter* and the rest to several other taxa. The increase in *Bacteroidetes* closely followed the phytoplankton summer bloom, as indicated both by chl *a* (Figure 2) and chloroplasts (Figure 3a).

*Actinobacteria* were present in small amounts throughout the study. Two more bacterial taxa, *Verrucomicrobia* and *Deltaproteobacteria*, were significant at the beginning of the time series, but eventually disappeared. Likewise, the two archaeal phyla *Thaumarchaeota* and *Euryarchaeota* were abundant in March and almost disappeared with the seasonal progression. The bloom of eukaryotic algae in June was reflected as a slight increase in the chloroplast reads towards the end of the study period (Figure 3a).

### Main iron transport genes

We searched for almost 250 unique genes grouped in 93 COG categories related to iron acquisition and siderophore synthesis (Supplementary Table S3). We pooled the genes into groups with similar functions (Table 1, see scheme in Supplementary Figure S3). Thus, the main transporters of iron were classified into: i) ferrous iron ABC transporters, ii) ferric iron siderophore outer membrane transporters (TonB dependent), and iii) ferric iron ABC transporters. These are the proteins destined to transfer ferrous iron across the cell membrane and ferric iron bound to siderophores across both membranes. Other categories were iv) divalent cation ABC transporters (transporting iron and other cations across the cell membrane), v) transporters of iron bound to dicitrate, vi) transporters of heme groups, vii) storage of iron as ferritin, and viii) regulators of iron metabolism. Finally, we also examined siderophore synthesis genes (see next section).

**Table 1.**
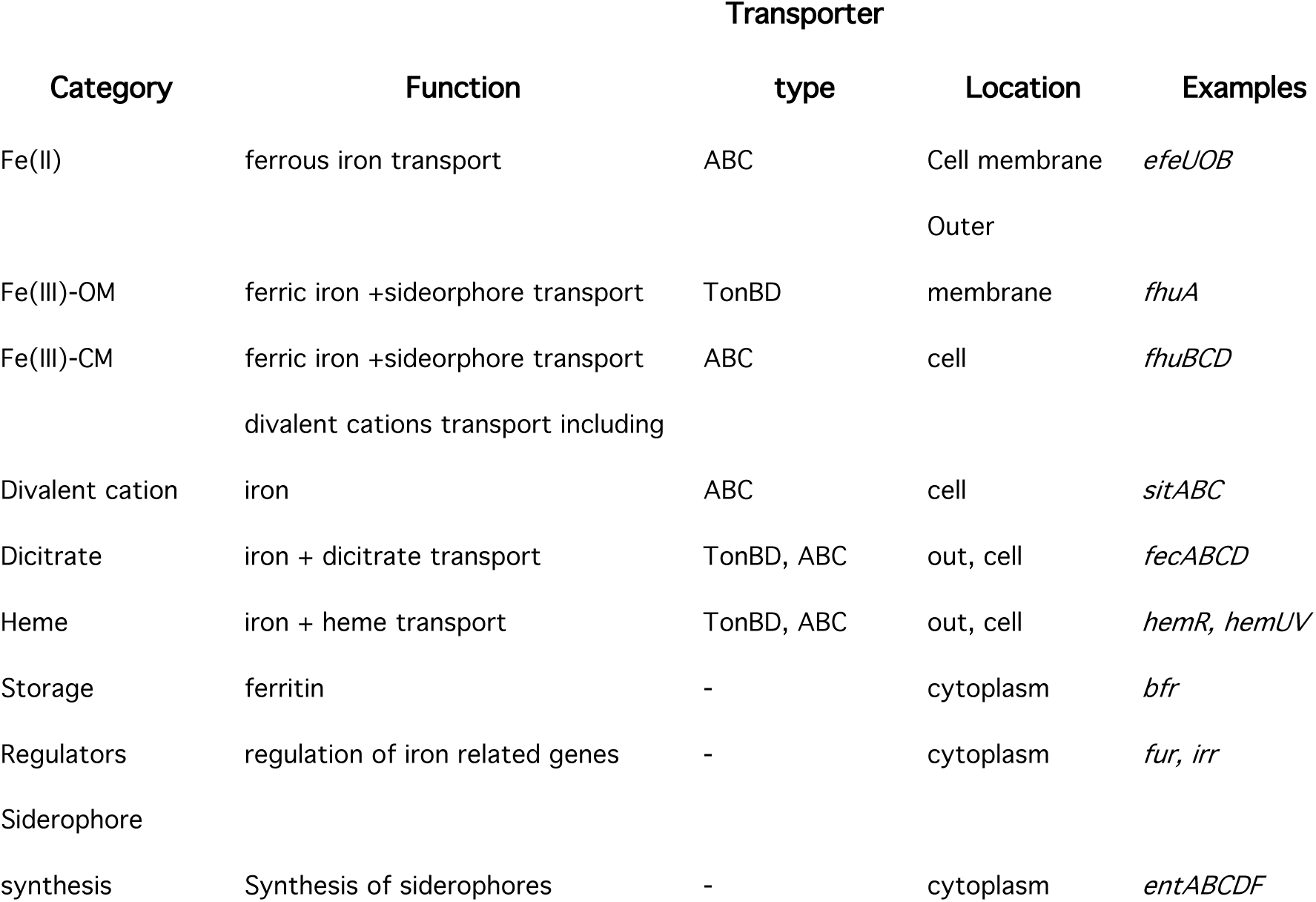
Categories of iron transport and siderophore synthesis genes used in the present paper.

Normalized abundance of iron related genes increased from a copy number (CN) of less than 5 in March to higher than 8 in late June (Figure 4a). Thus, the CN of genes devoted to iron acquisition almost doubled along the time series. Three of our categories, ferric iron-siderophore ABC transporters, storage, and siderophore synthesis did not change significantly. Two other, heme and divalent cation ABC transporters increased moderately, while ferric iron TonB dependent, dicitrate and ferrous iron ABC transporters increased substantially. Iron uptake regulation genes also increased and were those with highest CN throughout the study (between 2 and 4, Figure 4a).

**Figure 4.**
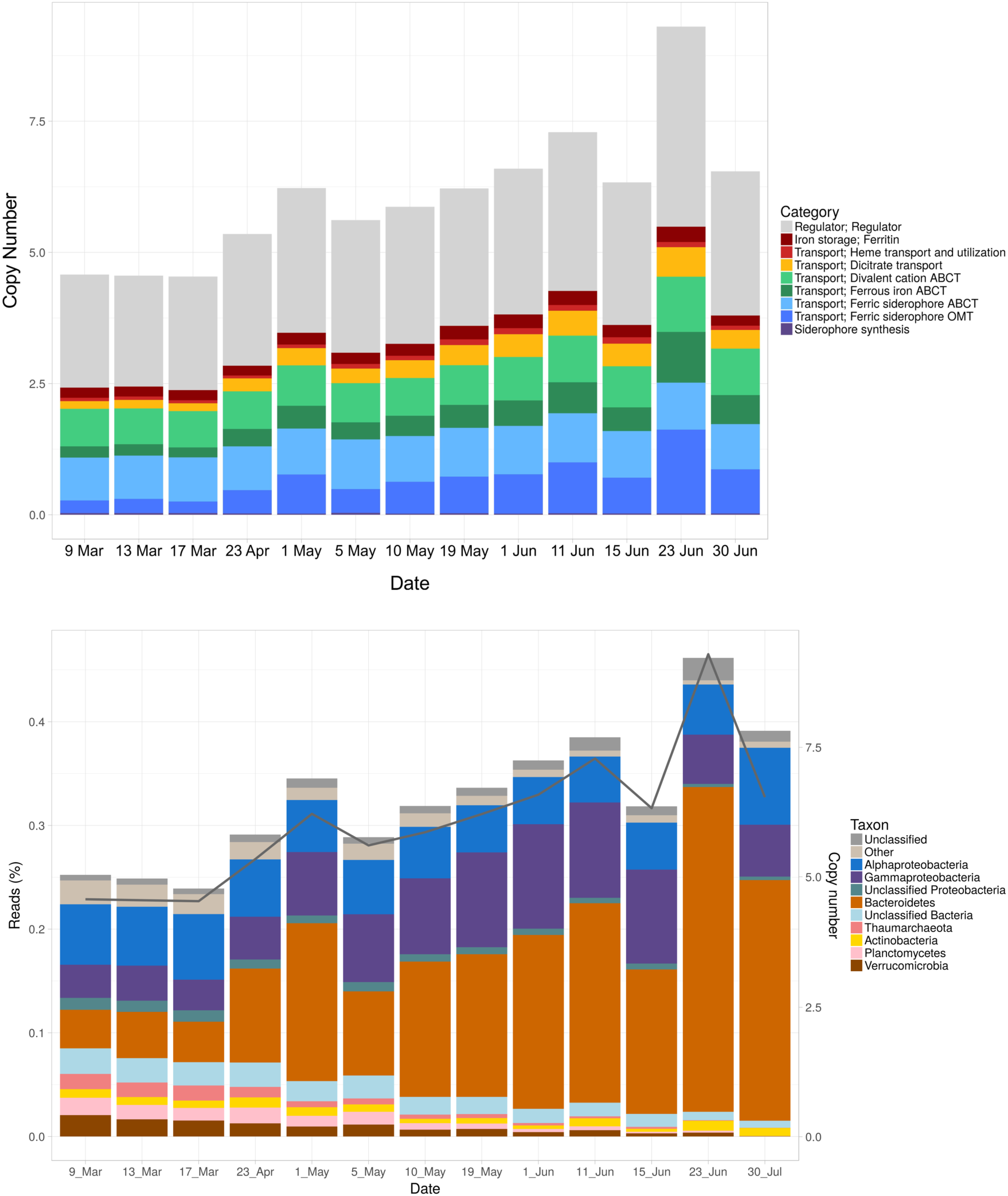
(a) Copy numbers of iron related genes through the study period in metagenomes. (b) Percent of the reads assigned to different taxonomic groups (bars, left hand scale) and copy numbers (line and right hand scale). **(b).** Taxonomy of iron related genes.

Globally, regulator genes accounted for about half of the copies in the category, ferric iron (TonB dependent and ABC) for a quarter and the remaining genes for another quarter.

The taxonomy of these genes generally corresponded to that of the community (compare Figs. 3a and 4b). In other words, the most abundant taxa were responsible for most of the iron related genes. There was only one obvious difference: while *Alphaproteobacteria* were always the first or the second group in abundance, they were always the third one when considering only the iron-related genes, with *Gammaproteobacteria* and *Bacteroidetes* alternating as those with the most genes. This underrepresentation was likely due to the fact that the most abundant member of the *Alphaproteobacteria* was the order *Pelagibacterales* (Figure 3b), that is known to lack all TonB dependent transporters and to have a very small percent of their genome devoted to iron acquisition genes (Cobo-Simón *et al*., in preparation).

### Ferric iron transport genes

There were large differences in abundance and taxonomy between the outer membrane and cell membrane transporters (Figure 5a and b). The TonB dependent outer membrane transporters increased from a CN of 0.3 in March to reach a maximum of 1.6 in late June. Most of this increase was due to *Bacteroidetes* (Figure 5a). In effect, this group accounted for 90% of the reads of these genes in late June. On the contrary, the inner membrane ABC transporters remained almost constant around a CN of 0.9. And the taxonomy was also completely different. In order of importance the main groups were *Alpha-*, *Gammaproteobacteria* and *Bacteoridetes* in the third place (Figure 5b).

**Figure 5.**
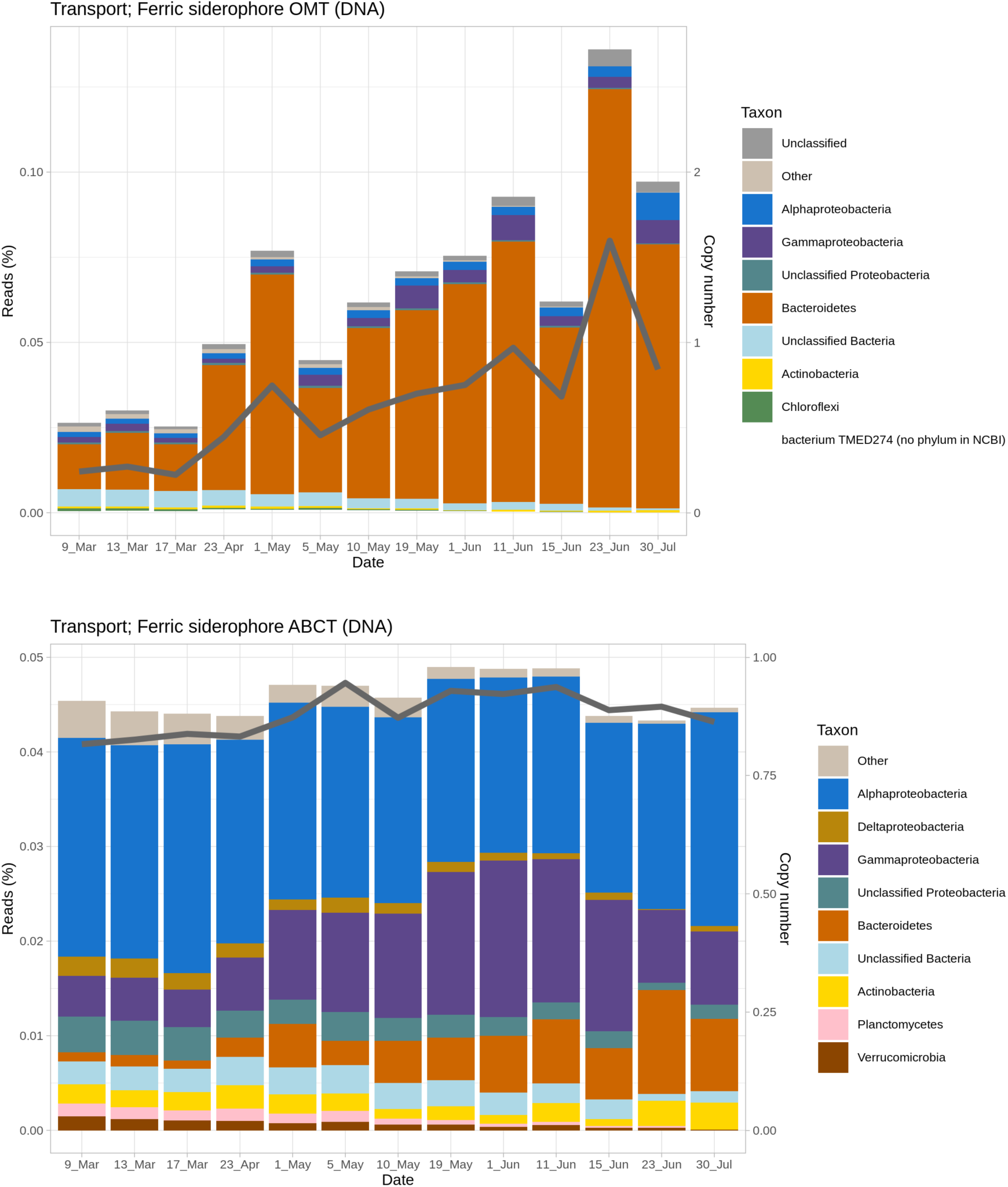
Taxonomic assignments and copy numbers of ferric iron transporter genes along the study: outer membrane ferric siderophore TonBD (above) and cell membrane ABC (below).

An alternative path to obtain ferric iron is incorporating it bound to dicitrate. The genes for this process increased in parallel to the outer membrane TonB dependent transporters, although at lower CNs (always below 0.6, Supplementary Figure S4a). In this case, *Gammaproteobacteria* and *Bacteroidetes* were responsible for most of the reads.

The last ferric iron transporter genes that we found were those devoted to transport heme bound iron (Supplementary Figure S4b). These genes also increased with time, mostly due to the *Gammproteobacteria*. *Bacteroidetes* did not have these genes. And towards the end of the study *Actinobacteria* and *Chlorophyta* contributed significantly. The CNs, however, were low, always below 0.15.

### Siderophore synthesis genes

The CN of siderophore synthesis genes oscillated slightly around 0.025 (Figure 6). These values are likely underestimated because we excluded all the reads of COG0318 (see M&M). Still, the CNs are very low compared to the genes to incorporate iron bound to siderophores. Thus, the effort dedicated to siderophore synthesis was moderate, and more or less constant. A portion of these genes could not be assigned to particular bacterial groups. But, of those that could be assigned, most belonged to *Proteobacteria*. Interestingly, *Beta*- and *Deltaproteobacteria* were significant contributors in March, while *Gamma*- and to a lesser extent *Alphaproteobacteria* were the main groups in June. A striking difference with previous groups of genes was that *Bacteroidetes* almost did not contribute anything to siderophore synthesis.

**Figure 6.**
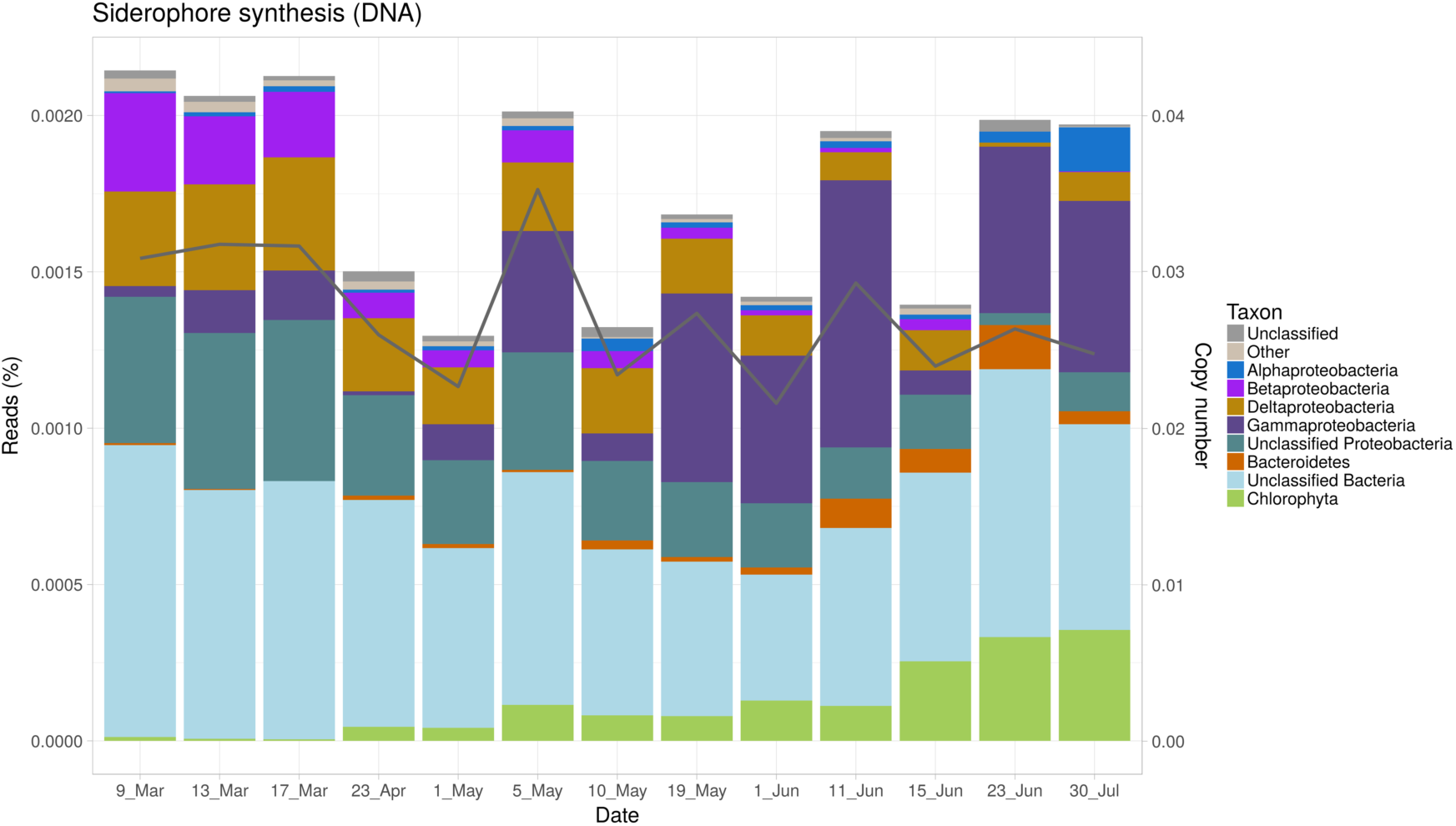
Taxonomic assignments (bars) and copy numbers (line and right scale) of siderophore synthesis genes.

With the criteria explained in M&M we could confidently state that pathways for eight siderophores were present: alcaligin, enterobactin, mycobactin, myxochelin, petrobactin, salmochelin, and staphyloferrins A and B, and that four were absent (pyochelin, pyoverdin, yersiniabactin, and amphibactin). Again, considering our criteria, the presence of the remaining studied siderophores could not be confirmed (Supplementary Table S4). All eight siderophore synthesis pathways identified as present were found in all the metagenomes except the earliest one on 9 March (Supplementary Table S5).

### Ferrous iron transport genes

Ferrous iron is able to diffuse through porins into the periplasmic space. There it can be transported to the cytoplasm by ABC transporters, either iron specific or able to transfer several divalent cations including ferrous iron. The ferrous iron specific ABC transporters increased during the study from a copy number of 0.25 to almost 1 in June (Figure 7a). Again, most of the increase was due to *Bacteroidetes* with *Gammaproteobacteria* having a constant contribution. Several other bacterial groups had significant contributions in March and disappeared along the study. These included *Actinobacteria*, *Planctomycetes*, *Verrucomicrobia* and *Gemmatimonadetes*.

**Figure 7.**
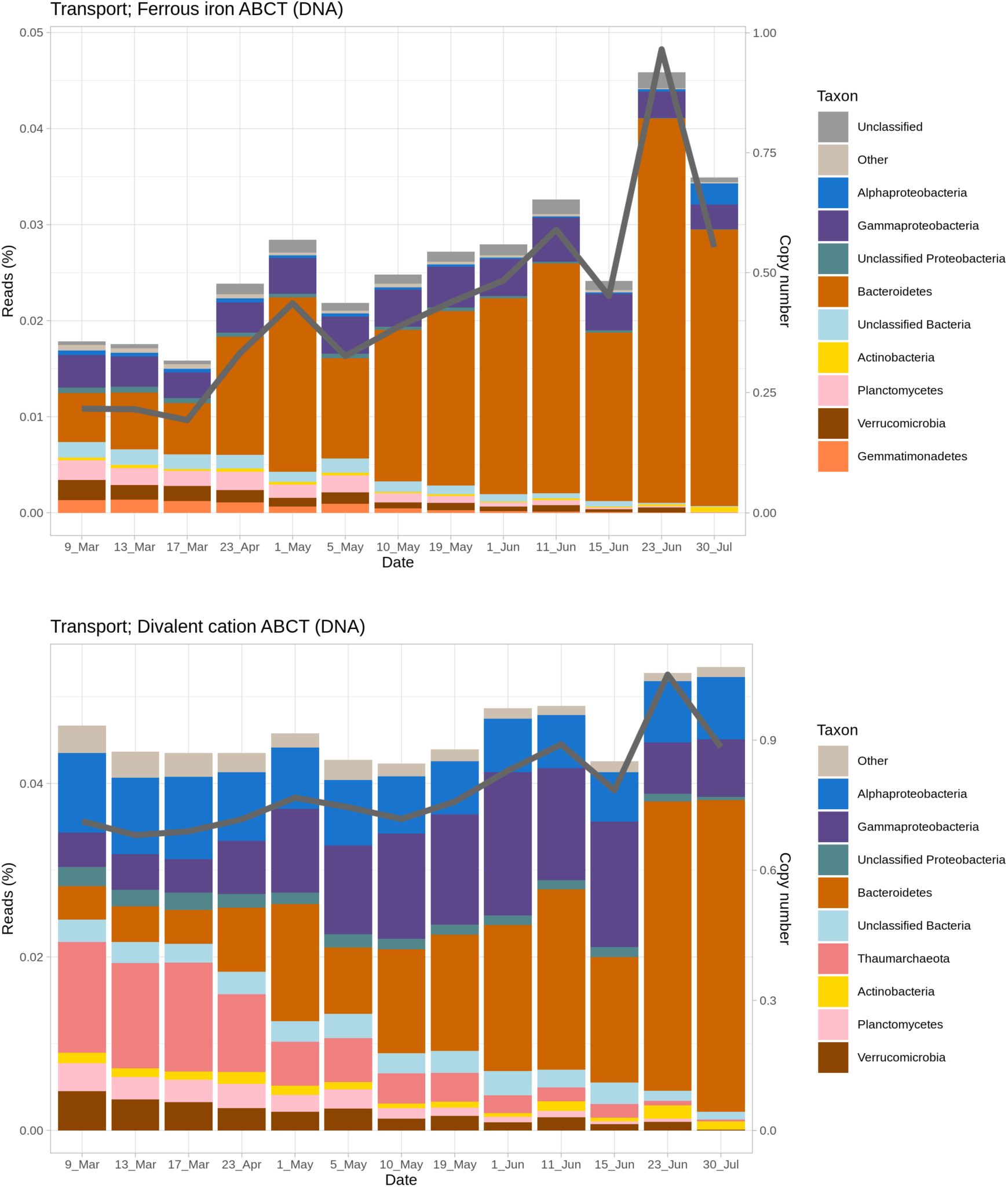
Taxonomic assignments and copy numbers of (a) ferrous iron ABC transporter genes and (b) divalent cation ABC transporter genes along the study.

The divalent cation ABC transporters, on the other hand, increased very slightly from a CN of 0.8 to 1.0 (Figure 7b). Interestingly, *Thaumarchaeota*, *Planctomycetes* and *Verrucomicrobia* accounted for half of the reads in March and decreased through spring. *Alphaproteobacteria* accounted for a constant proportion. *Gammaproteobacteria* increased until 15 June and then decreased again. Finally, *Bacteroidetes* had the largest increase becoming dominant in June (70% of the reads).

### Other iron related genes

We also looked for iron storage genes, mostly related to ferritin (Supplementary Figure S5a). There was only a moderate increase in CN, from 0.2 to 0.3. *Thaumarchaeota* were abundant in March, while *Gammaproteobacteria* and *Bacteroidetes* were abundant in June. Finally, genes involved in the regulation of iron metabolisms were always the most abundant ones (Supplementary Figure S5b), with CNs increasing slightly from 2.1 to 3.8 The three most abundant groups of bacteria were also responsible for most of the reads belonging to these genes.

### Gene expression

Taxonomic assignments of the metatranscriptomic reads are shown in Figs. S6 and S7. On several dates more than half of the reads were either unmapped or unclassified. As a result, the information they provide is limited. Thus, we have only used the metatrancriptomes to show that, considering the mapped and classified reads, expression was proportional to abundance of the taxa. Therefore, there were no differences among the different taxa in the level of expression.

## Discussion

### Community composition

The observed change in microbial composition from early spring to summer has previously been reported for the Amundsen Gulf (6), which is approximately 600 km from our study site. The winter community is generally dominated by *Alpha-* and *Gammaproteobacteria*. *Archaea* are important in winter and decrease towards spring and summer (33). Other minor but significant groups in winter are *Betaproteobacteria*, *Verrucomicrobia* and *Actinobacteria*. All of these groups typically decrease and even disappear in summer (8). In contrast, *Bacteroidetes* increase along the spring and become one of the dominant taxa. Predominance of the genus *Polaribacter* is known to be common in polar waters (34). This general structure of groups *Alpha*- and *Gammaproteobacteria* and *Bacteroidetes* dominating is fairly typical of the Arctic Ocean (8), and is supported by this study. This is in contrast to the usual community composition in temperate waters, where *Alphaproteobacteria* are generally dominant accounting for around 50% of the community, and are followed by *Gammaproteobacteria* and *Cyanobacteria*, while *Bacteroidetes* usually account for only 4% and *Betaproteobacteria* are barely detectable (8).

In general, both abundance and expression of genes involved in iron uptake was proportional to abundance of the organisms. *Gammaproteobacteria*, *Alphaproteobacteria* and *Bacteroidetes* were the most important. Debeljak et al. (35) studying heterotrophic bacterial assemblages around Kerguelen Islands, found that *Gammaproteobacteria* and *Bacteroidetes* were the main taxa in the uptake of iron-siderophore complexes, while *Alphaproteobacteria* had a more significant role in Fe(II) and Fe(III) uptake, just like we found. Likewise, Fourquez et al. (36), using microautoradiography, found that *Gammaproteobacteria* and *Bacteroidetes* were the main bacterial taxa active in iron uptake.

### Iron in the Arctic Ocean

There have been several studies determining iron concentrations in the Arctic Ocean. Most of these have been either in the Central Arctic close to the Eurasian coast ((37), (38), concentrations between 0.4 and 0.6 nM) or in the Chuckhi Sea and the Canadian Basin (39), (40), (41), (42). In the Bering Sea, Aguilar-Islas et al. (43) found iron to be limiting in the outer shelf not influenced by ice, but it was sufficient in the outer shelf influenced by ice. This was due to release of dissolved iron from the melting ice. To our knowledge, Colombo *et al*. (44) presented the only study in the Canadian Archipelago. Here, a higher concentration of iron was found than in the deep-water areas to the East and West, with surface concentrations between 0.40 and 1.91 nM above 40 m and between 1.10 and 1.15 between 70 and 100 m. They also noted the well-known inverse correlation between iron concentration and salinity, due to the known influence that freshwater river discharge and ice melt have on enriching sea water in iron. Therefore, iron concentrations in the area are much higher than in most of the oceans. Moreover, N:P ratios in the ice-water interface at our study site suggested nitrogen limitation (21). Thus, iron was not expected to be limiting in these waters. Despite this, iron is an essential nutrient for all microorganisms and it requires special proteins to acquire it.

While iron in ocean water could not be directly sampled in this study, our metagenomic data showed presence of iron acquisition genes in all three dominant bacterial taxa, with copy numbers larger than 1 in some cases.

Unlike the Arctic Ocean, the Southern Ocean is one of the main HNLC areas in the world, where phytoplankton growth is limited by iron. However, there is considerable variability. Thus, in studies of the waters around Kerguelen Islands, Obernosterer *et al*. (45) could compare HNLC waters on the windward side of the islands with naturally iron enriched waters on the leeward side. These authors found that heterotrophic bacteria were limited by iron in both regions. Debeljak *et al*. (35) analyzed the expression of several iron acquisition genes in the same area and found that transcripts for Fe(II) and Fe(III) uptake genes were found in higher proportions at the HNLC stations as compared to the Fe-fertilized sites. In contrast, no such pattern was observed for siderophore-uptake (35). However, when they analyzed the taxa specific expression, siderophore uptake transcripts were higher in the HNLC station than in the iron enriched zones. In principle, we should expect in the Arctic Ocean a situation similar to that in the iron enriched zone. That is, an abundance of iron uptake genes with those for siderophores outnumbering those for Fe(II) and Fe(III) and, in effect, this is what we found. Therefore, it seems a general feature of marine bacterial assemblages that iron acquisition is a major need of bacteria even under non limiting conditions.

### Siderophores

In the aerobic ocean, iron is predominantly in the Fe(III) oxidation state, which has a very low solubility and is therefore hard to obtain. One of the most common ways for microorganisms to thus acquire ferric iron is to synthesize siderophores, that can be carried out by many marine microorganisms (11). The chemistry of siderophores in marine microorganisms has been reviewed (46), (47). These studies involve pure culture isolates of marine bacteria in which the siderophores are produced in large enough quantities to be characterized chemically. Most of these belong to *Gammaproteobacteria* genera *Halomonas*, *Vibrio*, *Marinobacter*, *Ochrobacter*, *Pseudoalteromonas*, *Shewanella*, as well as cyanobacteria such as *Synechoccocus* (15).

Concentrations of siderophores have been measured in a vertical profile at ALOHA station off Hawaii (48), transects from the coast offshore in the southeast Pacific (49), the California Current System (50), and along a latitudinal transect in the oligotrophic Atlantic Ocean (51). In all these studies, ferrioxamines were found in surface waters, while amphibactins were typically more abundant in deep waters and in high nutrient low chlorophyll zones. Extrapolating this distribution to the costal Arctic, we would expect genes for ferrioxamines but not those of amphibactins in our 60 m deep study region. Actually, we found the whole set of genes for ferrioxamines B and E, which would be in agreement with the previous studies. However, none of these genes was unique to these pathways and, therefore, we cannot assert whether these siderophores were present in our samples or not.

The pathways for synthesis of amphibactins have not been as well characterized. However, the pathway in *Vibrio neptunius*, a pathogen of bivalve mollusks has been determined (52). In our metagenomes we found all the genes for this pathway but one. Moreover, all the genes found were common to other siderophore synthesis pathways. According to our conservative criteria, amphibactin synthesis pathways were not present in our samples. This would agree with the absence of amphibactins from surface ocean waters. Amphibactins have a polar head with the function of binding iron linked to a fatty acid tail that may be anchored in the outer cell membranes, thus avoiding the dispersion of the siderophore beyond the reach of the producing organism (53) (15).

This is a way to avoid utilization of siderophores by alien microorganisms and would seem an adaptive strategy in the oceans. However, as discussed, they do not seem to be abundant in surface marine waters and we did not find the complete pathway in our study.

### Siderophores in Proteobacteria and Bacteroidetes

*Proteobacteria* is the bacterial phylum in which siderophore synthesis and uptake has been most studied, particularly concerning pathogenic *Gammproteobacteria* (54), (55). In contrast, experimental evidence of siderophore synthesis in *Bacteroidetes* is scarce. Some pathogenic *Flavobacteria* can synthesize siderophores with genes in a plasmid (56). Delmont *et al*. (57) found siderophore synthesis genes in three *Bacteroidetes* metagenomic assembled genomes (MAGs), including two *Polaribacter* and one *Cryomorphaceae,* in a bloom in Antarctica. The *Proteobacteria* MAGs they found did not have these genes. Guan *et al*. (16) found a ferrichrome (*fhuA*) receptor in the fish pathogen *Flavobacterium columnare* that was shown to also synthesize siderophores with the chrome azurol assay. Chen *et al*. (47) reviewed siderophores from marine bacteria and found over 80 of all the known siderophore families. Most siderophores were synthesized by *Gammaproteobacteria* and a few by *Alpha*- and *Betaproteobacteria* and *Actinomycetes*. Only one siderophore was synthesized by a *Bacteroidetes*. This was bisucaberin B made by *Tenacibaculum mesophilum* isolated from a tropical sponge. Fujita *et al*. (58) claimed this was the first siderophore made by a *Bacteroidetes*. In *Allivibrio salmonicida* (*Gammaproteobacteria*) the gene cluster for synthesis and uptake of this siderophore is flanked by transposases. This suggests that this cluster may have been acquired by lateral gene transfer. The same might have occurred in *Tenacibaculum*. Bisucaberin was one of the siderophores that might be synthesized in the Arctic, but the reads did not belong to *Bacteroidetes*.

On the other hand, there is evidence of *Bacteroidetes* being able to use siderophores made by other bacteria. D’Onofrio *et al*. (17) found that a *Maribacter polysiphoniae* (a member of the family *Flavobacteriaceae*) could grow on solid agar when supplemented with a siderophore from a producer bacterium such as *Microccocus luteus* isolated from the same sediment environment. Zhu *et al*. (18) showed that *Bacteroides thetaiotaomicron* could use enterobactin and salmochellin from *E. coli* and *Pseudomonas* in the intestine with *xusABC* genes. Our results are consistent with this model of *Bacteroidetes* contributing very little to synthesis but considerably to uptake of siderophores.

### Iron-siderophore complexes as common goods

In our Arctic study, the time series followed the development of the seasonal phytoplankton bloom (5), which is triggered by the increasing intensity and duration of solar radiation available (59). In addition to the increased growth of primary producers as ice melts, large amounts of polysaccharides made by ice-algae may be released in the water column (60). For this reason, *Bacteroidetes* are known to grow better when accompanying algal blooms and have a large array of outer cell membrane proteins to take up polysaccharides from them (61), (62). This is likely the main factor in the dominance by *Polaribacter* within this study. However, given the substantial amounts of genes for siderophore uptake and their active expression, it seems that iron was a desirable commodity. Our data on the siderophore synthesis and uptake genes are consistent with competition for siderophores between *Gammaproteobacteria* and *Bacteroidetes*. Both taxa dedicated most iron acquisition genes to the uptake of iron- siderophores. *Bacteroidetes* seemed to act as cheaters while *Gammaproteobacteria* were synthesizing them. It seems likely that this characteristic, among others, may have helped *Bacteroidetes* to become the most abundant group at the end of the study. Our results confirm the idea of siderophores as public goods, highlight the fact that iron-siderophores are competitively sought after even in environments not limited by iron, and open a fascinating avenue of research for future Arctic cruises.

## Supporting information

Suppelemntary tables

## Author contributions (not necessary in ISME journal)

Conceived and designed the work: C.J. Mundy, CPA.

Acquired and processed data: C.J. Mundy, K. Campbell, Marta Royo-Llonch, Vanessa Balagué, P. Sánchez, L. Macías, F, Puente-Sánchez, J. Tamames.

Drafted the manuscript: L. Macías, F. Puente-Sánchez, CPA.

All authors contributed to discussion of the results, finalization and approval of the manuscript.

## Acknowledgements

Sampling in Dease Strait was funded by an NSERC Discovery and Northern Research Supplement Grant to C.J.M., and in-kind support from the Canadian High Arctic Research Station (CHARS). A. Delaforge assisted greatly in data collection and sample processing. Sequencing and data analysis were funded by grant PID2019-110011RB-C33 from the Agencia Estatal de Investigación of the Spanish Ministerio de Economía y Competitividad to J.T. and C.P.-A. Initial computational analyses were carried out at the Marine Bioinformatics Core facility MARBITS at ICM-CSIC Barcelona, as well as nucleic acid extractions. Most computation and analyses were carried out at CNB.

