## Supplementary figures for "Bacterioplankton taxa compete for iron along the early spring-summer transition in the Arctic Ocean"

Competition for iron along the early spring-summer transition in Dease Strait, of the Canadian Artic

**Supplementary Tables**

**Supplementary Tables supplied as a single Excel file**

**Supplementary Table 1**. Number of reads and mapped reads for each sample.

**Supplementary Table 2**. Details of the assembly.

**Supplementary Table 3**. Genes and COGs explored in the present work.

**Supplementary Table 4**. Siderophores with complete pathways (green) with incomplete pathways (orange) and with complete pathways but without unique genes.

**Supplementary Table 5**. Copy numbers of all the COGs for each date. Metagenomes are shown in pink and metatranscriptomes in blue. COGs in green are present in two or more pathways. Results in yellow show positive values.

**Supplementary Figures**

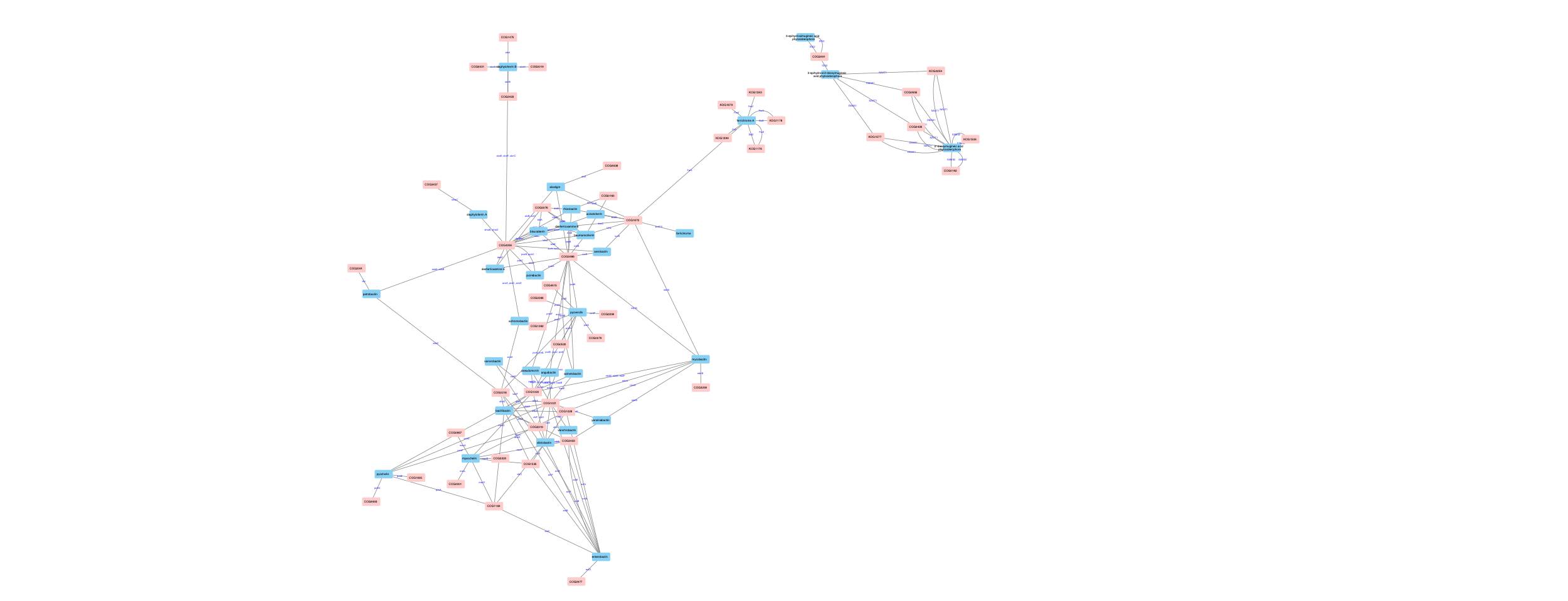

**Supplementary Figure 1.** Network showing COGs and siderophores. **See Cytoscape file for a good resolution, interactive image**.

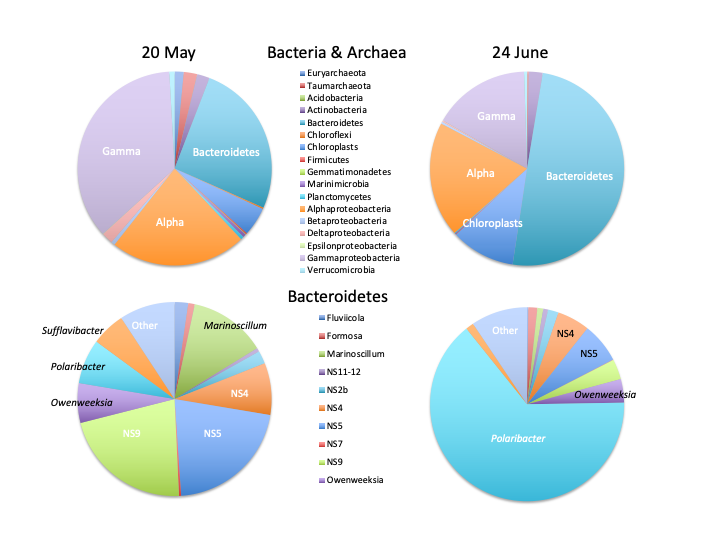

**Supplementary Figure 2.** Relative abundance of Bacteria and Archaea (pie charts above) and *Bacteroidetes* (pie charts below) on two dates along the winter to summer transition.

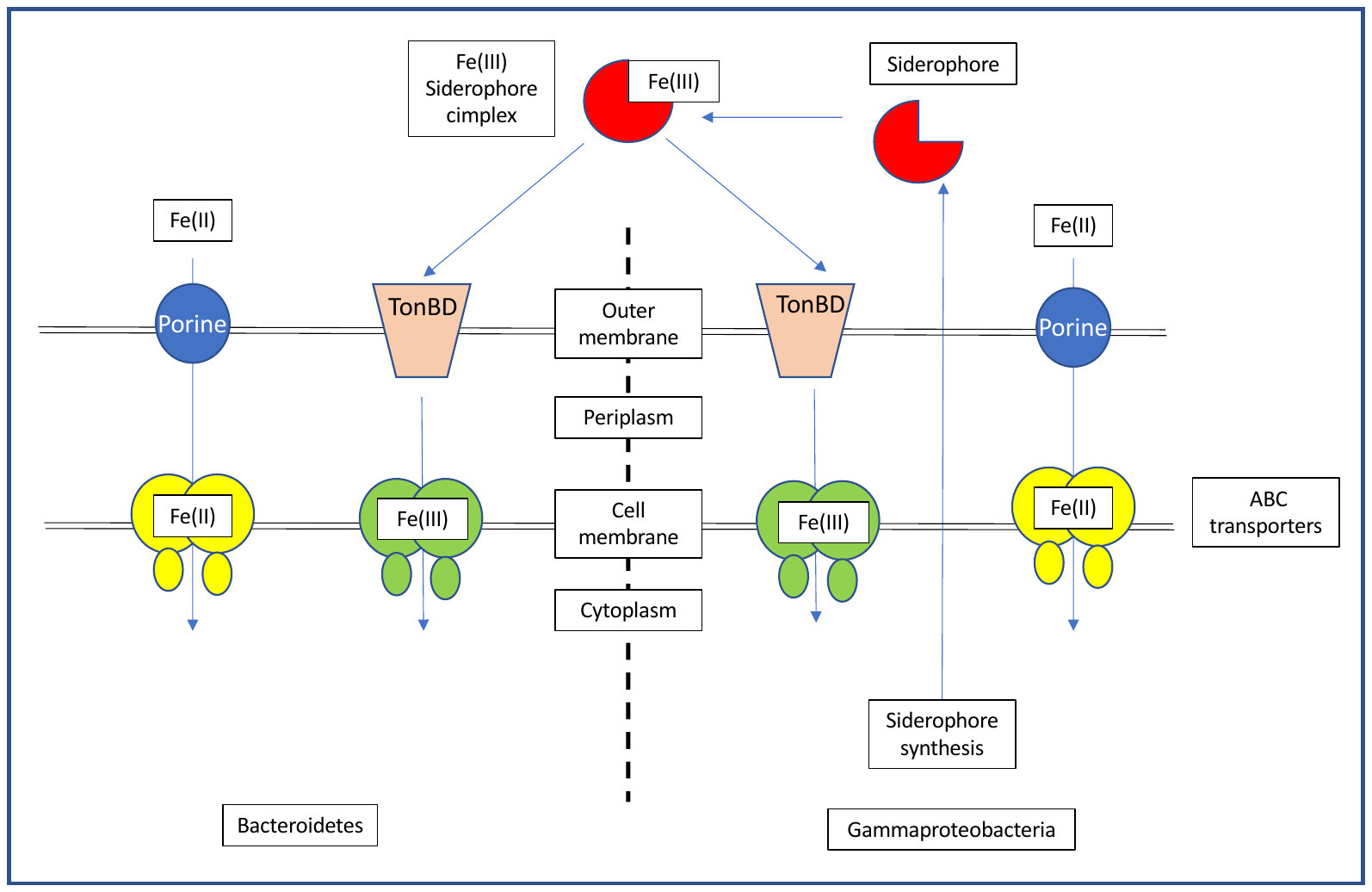
**Supplementary Figure 3.** Simplified scheme of iron uptake in *Bacteroidetes* (left) and *Gammaproteobacteria* (right). Fe(II) crosses the outer membrane through porines (blue) and the cell membrane through both Fe(II) specific and divalent cation generalist ABC transporters (yellow). Fe(III) requires binding to siderophores (red) to remain in suspension. These complexes are taken up by specific TonB dependent transporters (orange) in the outer membrane and cross the cell membrane through specific ABC transporters (green). Siderophores require a complex series of enzymes to be synthesized and excreted.

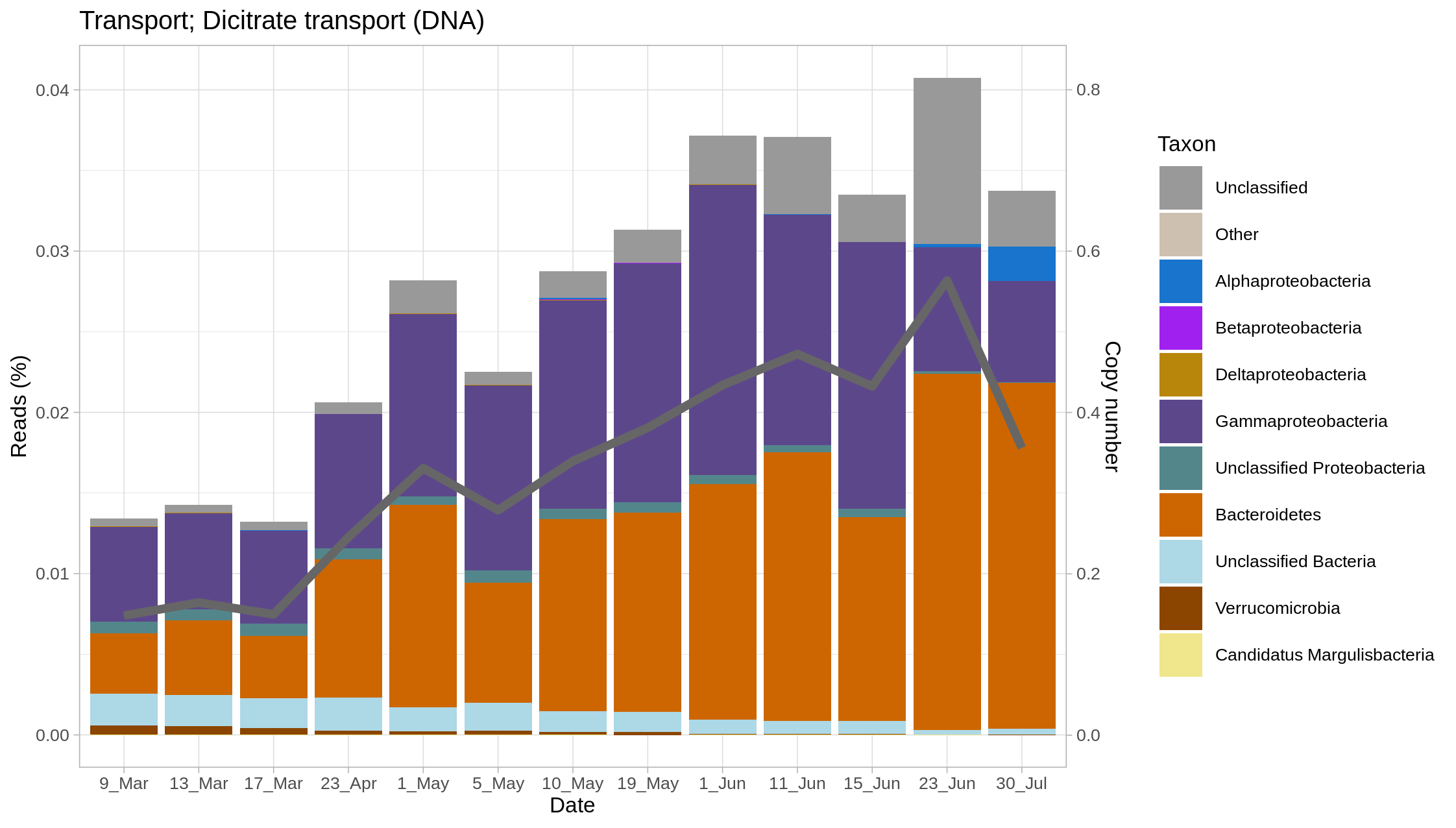

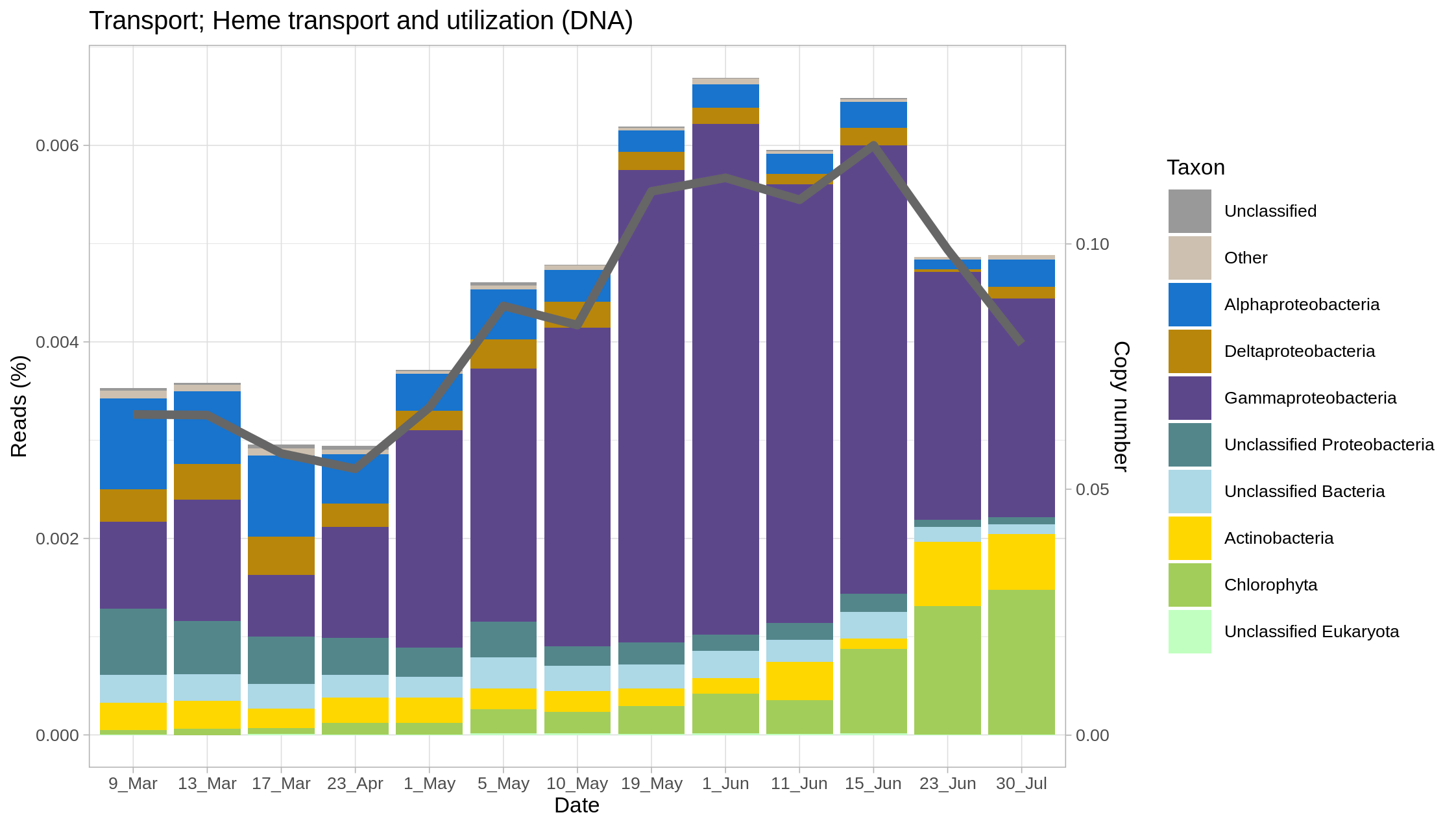
**Supplementary Figure 4.** Percentage of reads assigned to different taxa (bars, left hand scale) and copy numbers (line and right-hand scale) for dicitrate transporter genes (a) and heme transport genes (b) in metagenomes along the study.

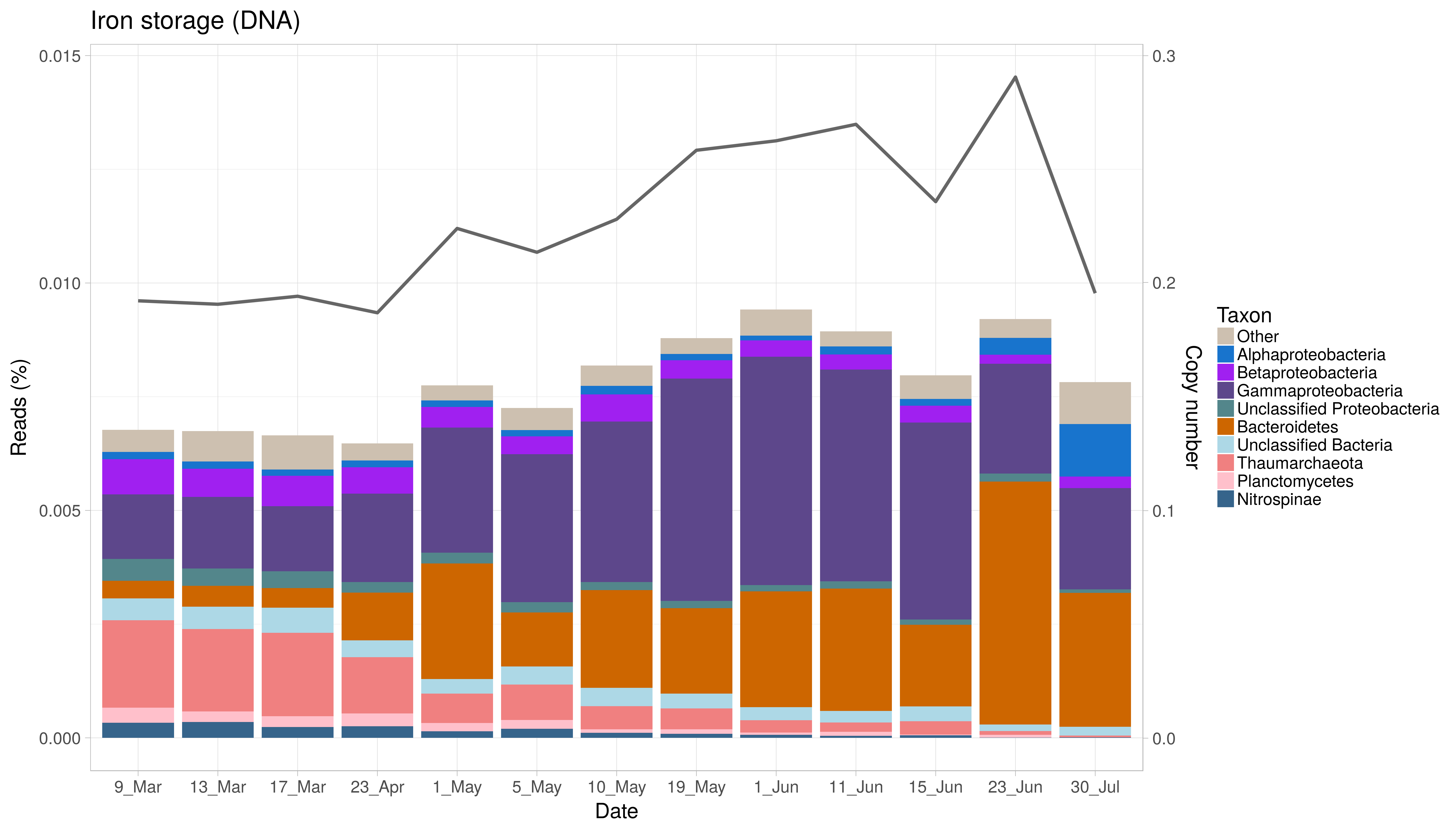

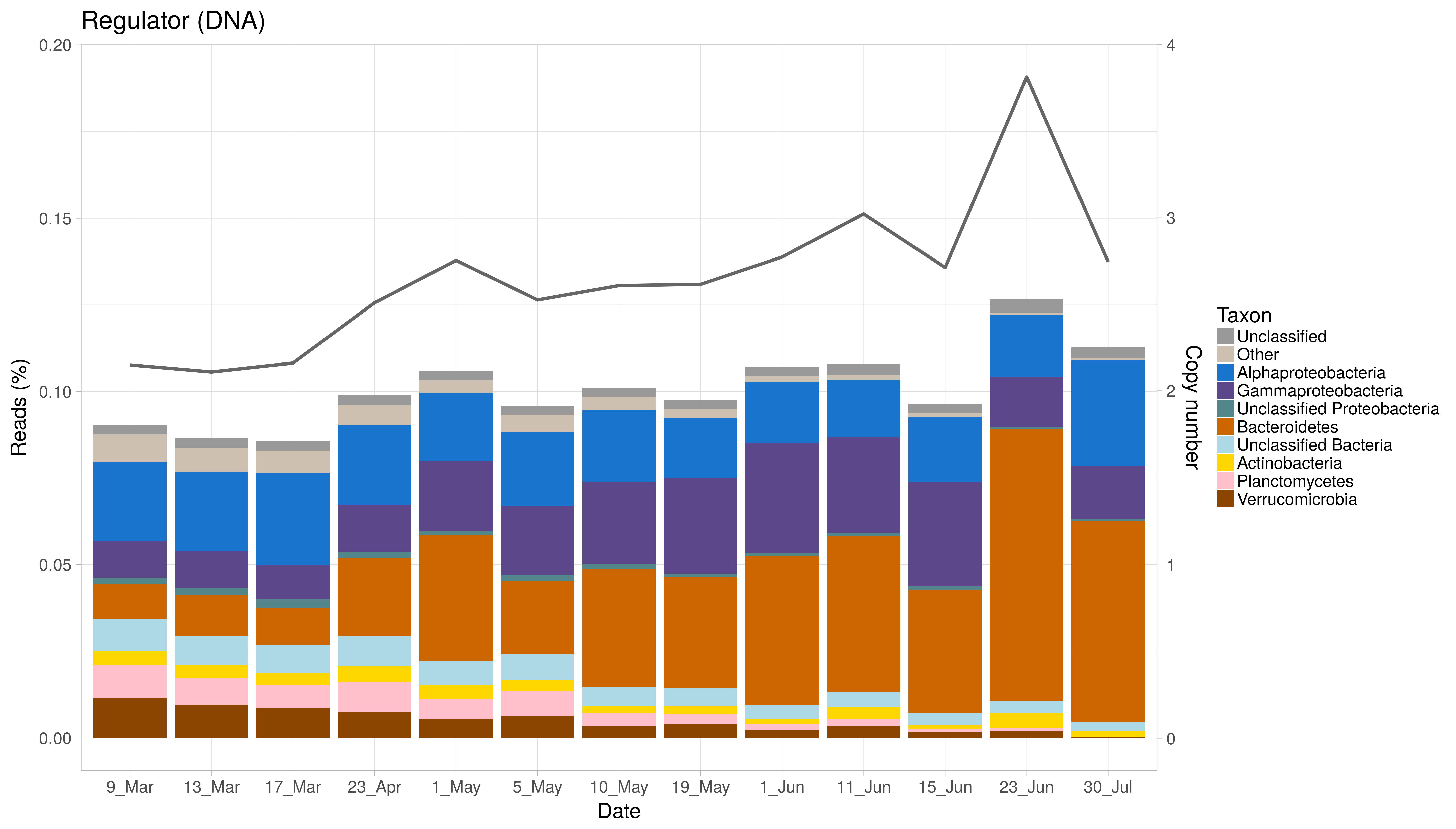
**Supplementary Figure 5.** Percentage of reads assigned to different taxa (bars, left hand scale) and copy numbers (line and right-hand scale) for iron storage genes (a) and iron regulation genes (b) in metagenomes along the study.

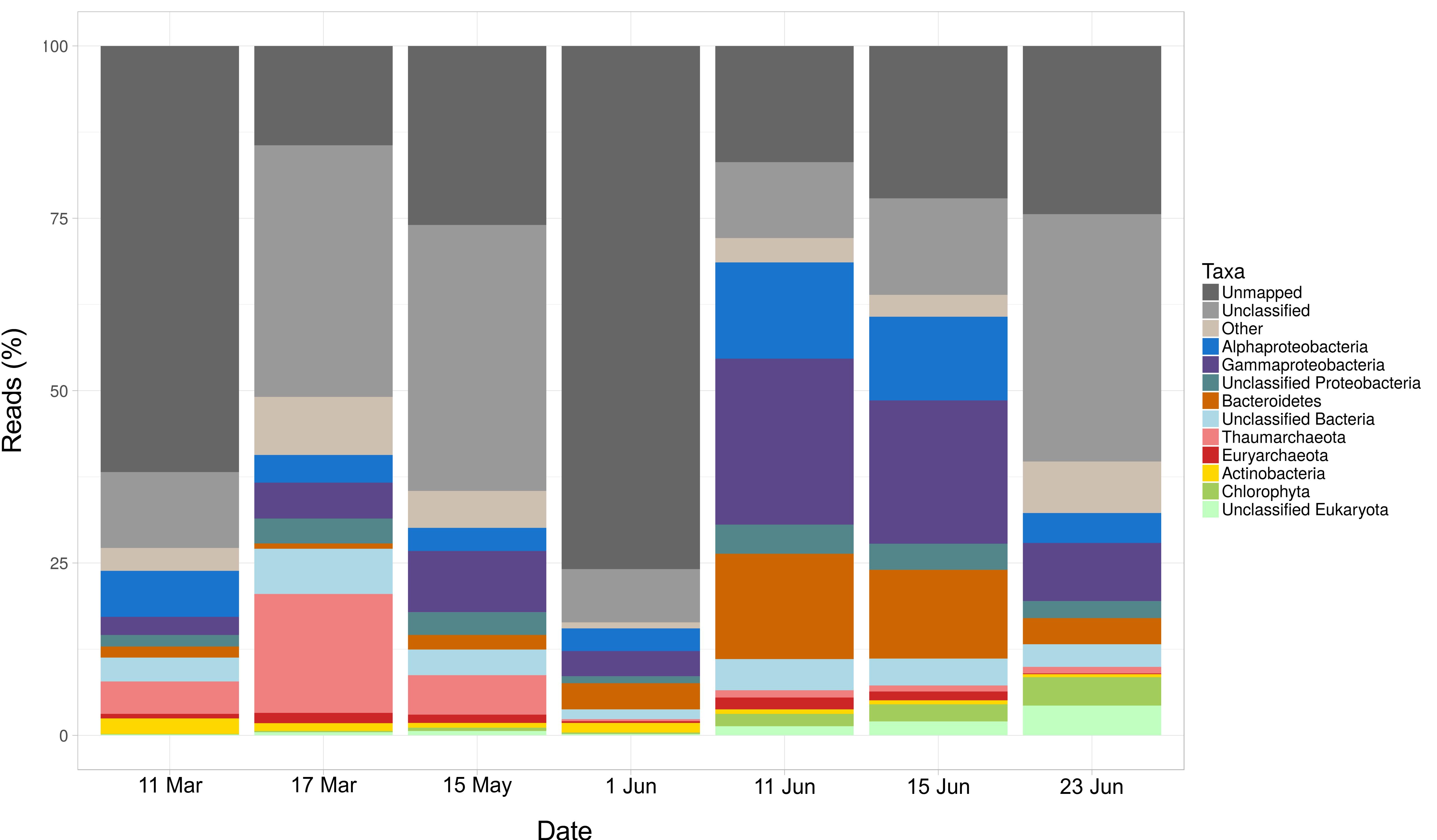

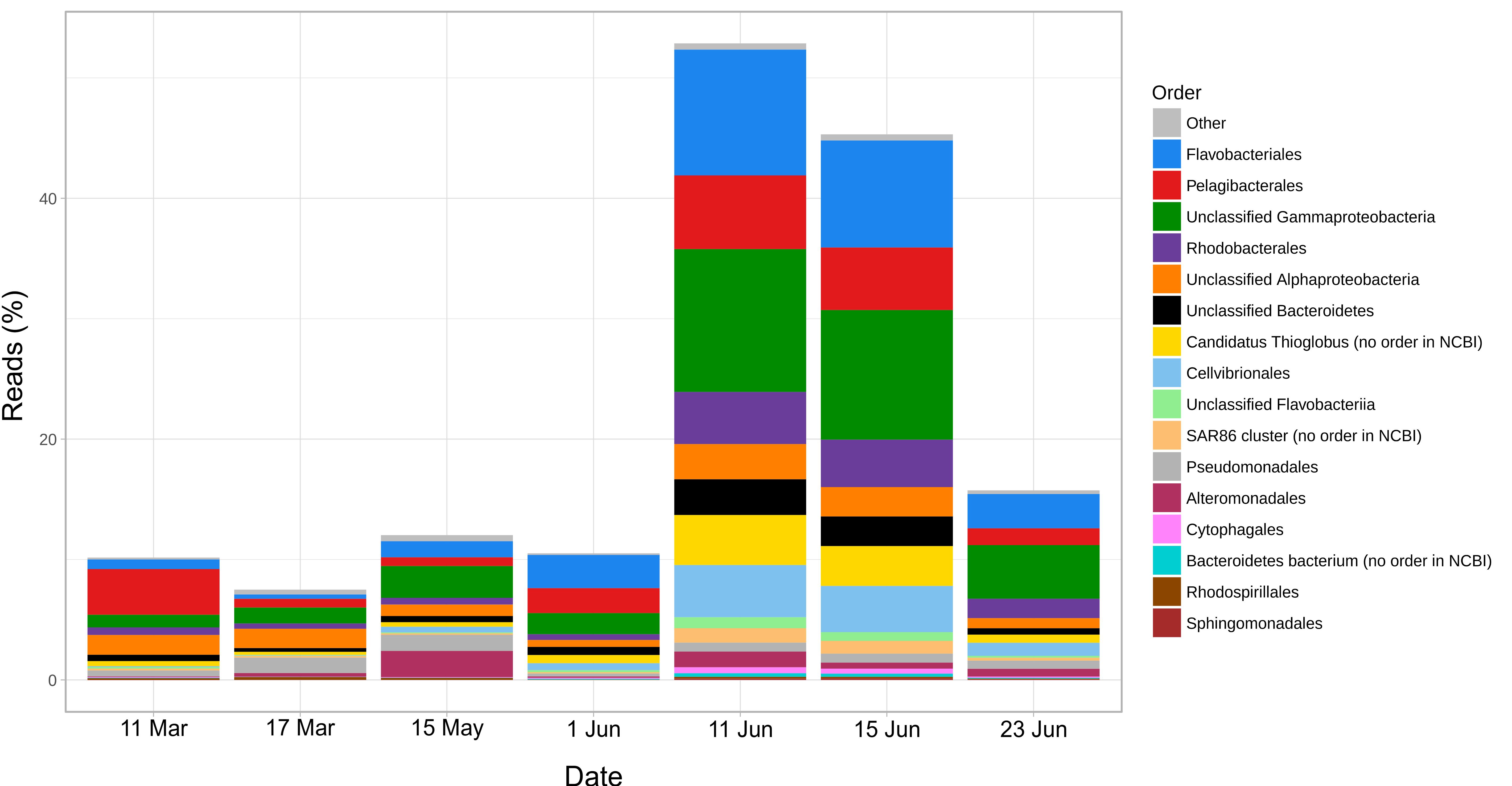
**Supplementary Figure 6.** Percentage of reads in metatranscriptomes at two different taxonomic levels: phyla and Proteobacteria classes (above) and orders (below).

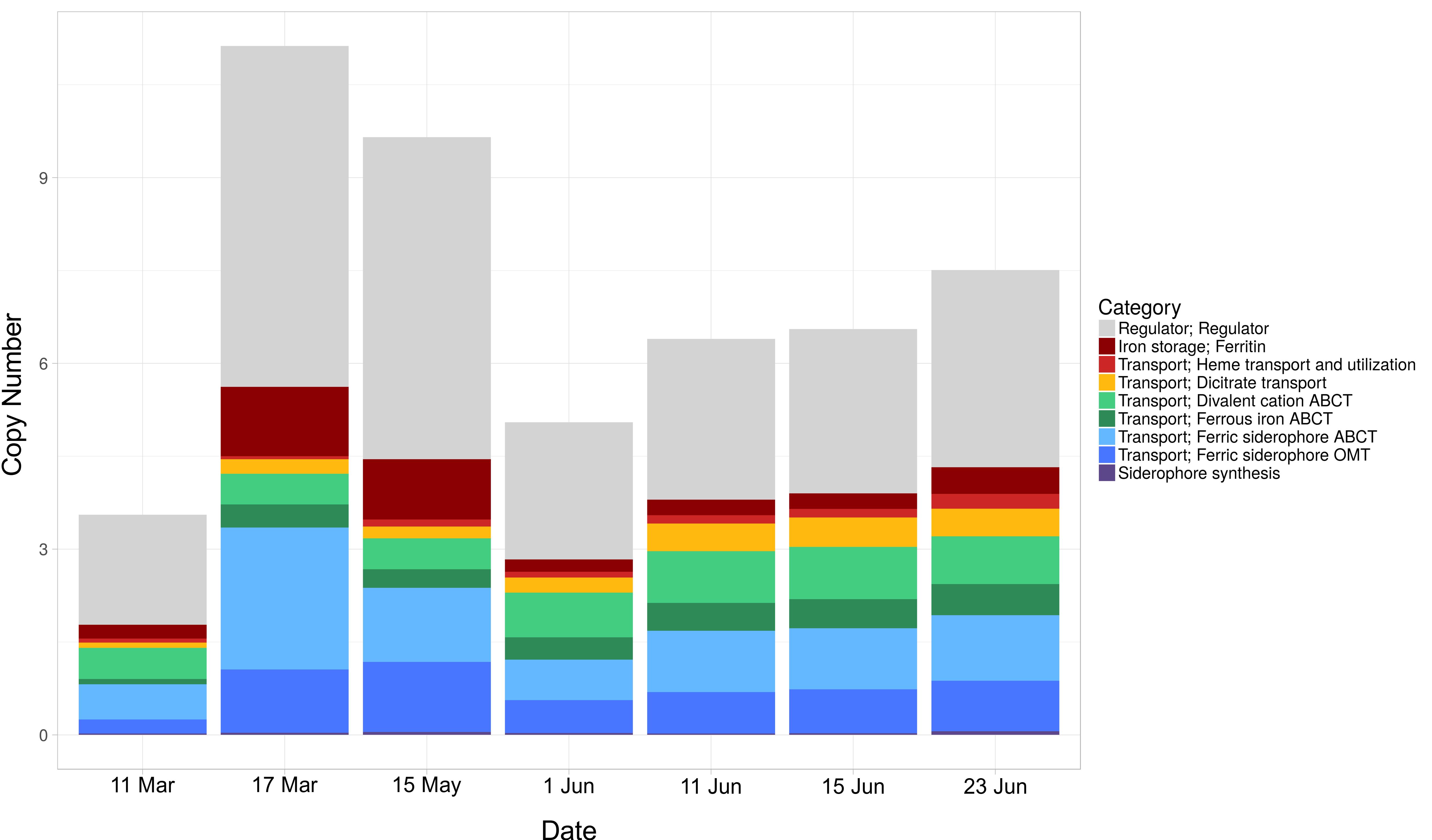
**Supplementary Figure 7.** Copy numbers of iron related genes through the study period in metatranscriptomes.
